# *k*-mer Distributions of Aminoacid Sequences are Optimised Across the Proteome

**DOI:** 10.1101/190280

**Authors:** A.A. Morozov

## Abstract

*k*-mer based methods are widely utilized for the analysis of nucleotide sequences and were successfully applied to proteins in several works. However, the reasons for the species-specificity of aminoacid *k*-mer distributions are unknown. In this work I show that performance of these methods is not only due to orthology between k-mers in different proteomes, which implies the existence of some factors optimizing k-mer distributions of proteins in a species-specific manner. Whatever these factors could be, they are affecting most if not all proteins and are more pronounced in structurally organized regions.

## Introduction

k-mer based methods are widely used in metagenomic studies because of their relatively low computational cost compared to aligning reads to refence database. The exact algorithms vary between implementations [1-3], but the idea is that *k*-mer spectra (or distributions) of phylogenetically close taxa are more similar to each other than they are to those of more distant groups. There is a plenty of empirical data to support this notion. The above-mentioned metagenomic approaches perform rather well on both simulated and real datasets, and *k*-mer based distance metrics have been used to reconstruct large-scale phylogenomic trees which were consistent with trees produced by more orthodox methods [4].

Most of the *k*-mer-related work in bioinformatics was performed on nucleotide sequences, but there is nothing inherently DNA-specific in this kind of analysis. There are works that have translated *k*-mer based methods, initially designed for DNA, to proteomics. Using a distance metric based on relative frequencies of *k*-mers, [5] have reconstructed a phylogenetic tree of 109 different organisms from all major taxa. The topology of this tree does not contradict results produced by other methods. In more recent work [6], a tree of approx. 900 bacteria with some eukaryotic outgroups was built using a different distance metric, again pretty consistent with the consensus on bacterial evolution. A recent metagenomic classifier named Kaiju [1] leverages protein conservativity to classify sequences that don't have any close relatives in the reference database. Thus, there is no question of whether *k*-mer distribution in aminoacid sequences is species-specific or whether the divergence of these distributions correlates with evolutionary distances. However, there is no answer to *why* it does.

The most common explanation relies on the orthology between *k*-mers in query sequence and database. When the classifier is concerned with orthologous sequences, as eg in case of classifying SSU RNA reads via RDP classifier [7], with sufficient value of *k* the chance of identical *k*-mers appearing in non-homologous parts of sequences by random coincidence is negligible. Somewhat similarly, protein-level metagenomic classification in Kaiju relies on finding MEMs (maximum exact matches) and extending them to inexact shared *k*-mers. While not stated explicitly, the phylogenetic importance of shared subsequences is also based on the orthology assumption. However, performance of *k*-mer-based classifiers and distance metrics on divergent bacterial proteomes with relatively few shared genes suggests there may be more to *k*-mer distribution than MEMs. In this work I show that this specificity holds even in the complete absence of the orthology.

## Results and Discussion

Performance of the naïve bayesian classifier on CEGMA dataset is shown at fig.1. In practically all cases this classifier performs better than random, and with optimal *k* of 5-7 more than 50% of sequences are assigned correctly. There is no possible orthology between sequences from the same species' training and test sets. In fact, there is a risk that a protein from test set has an ortholog in the *wrong* species' training set. *k*-mer distribution specificity persists even despite the lack of orthology, which suggests that it is formed by species-specific factors on the proteomic scale, rather than solely by the requirements of a particular protein family. Expanding the dataset to the entire proteomes leads to precision skyrocketing to almost 100%. Although some part of the precision increase can be explained by the presence of recently duplicated paralogs and isoforms, it still suggests that most, if not all, proteins are affected by these factors.

**Figure 1.**
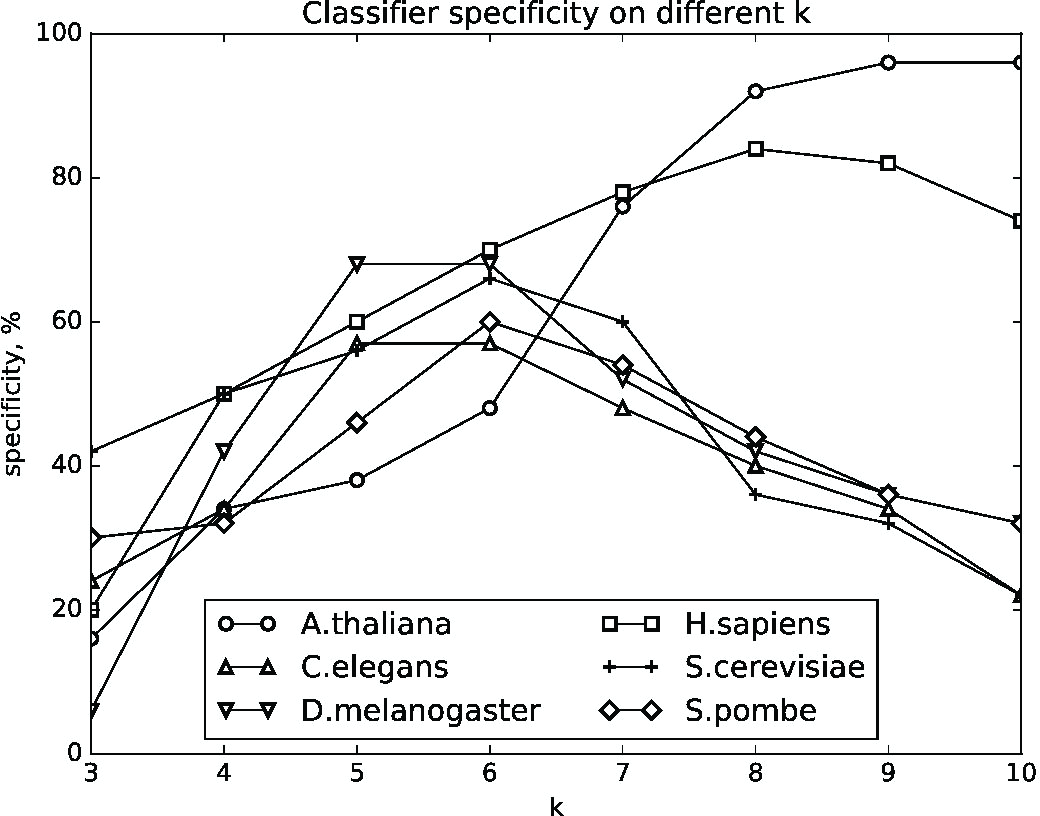
Specificity of naive Bayesian classifier on CEGMA dataset under different values of *k*.

To study the effect of these factors on a finer scale, we have built *k*-mer distributions for protein features from the complete proteomes of the same species according to UNIPROT annotations. Distances between the *k*-mer distribution of the feature in a particular species and the summary distribution for this feature across the entire dataset were calculated. The higher this distance, the more different these features in one organism are (on average) from their counterparts from other species, which allows to use them as a proxy for the species-specificity of *k*-mer distribution in protein fragments. As only structural features and entire domains have both average length and feature counts sufficient for a reliable estimation of *k*-mer distribution, various binding sites and signal peptides are omitted. Box-plots of these distances among different species are shown at fig. 2.

**Figure 2.**
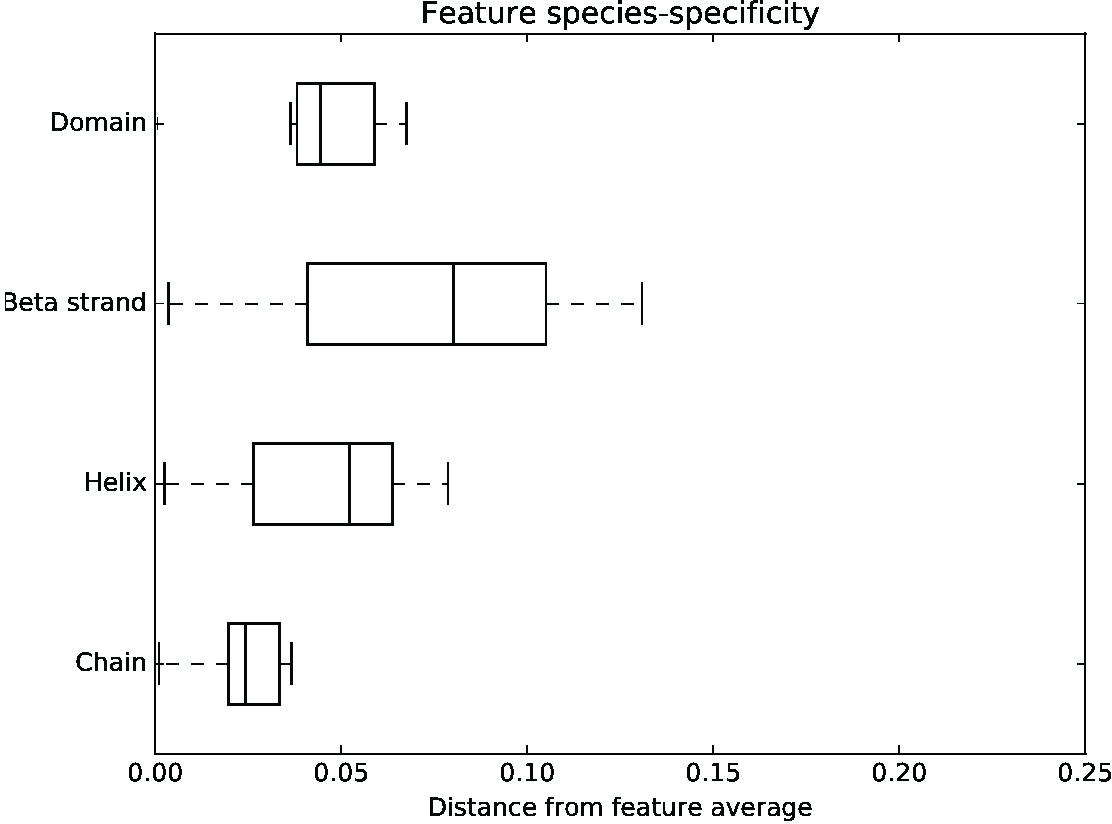
Species-specificity of *k*-mer distribution on different features across six proteomes. “Chain” feature represents protein sequence as a whole.

For all structurally organised elements (*ie* helices and beta-strands) *k*-mer distributions are more species-specific than they are for protein sequences as a whole (fig. 2), which means that pressure for *k*-mer adaptation is greater in this regions. The same is true for functional domains, whose *k*-mer distributions are optimised above protein-average level. This is strikingly similar to codon usage adaptation on DNA level, where the use of different codons is regulating kinetics of translation and folding. In particular, quickly translating high-frequency codons are common in alpha helices, while rare, slower ones are more likely to be found in random coils [11]. Several mechanisms can be proposed to explain this specificity on protein level. It's possible that the evolutionary advantage or disadvantage of particular *k*-mers is related to protein creation specifics, *eg* quicker and more efficient folding of optimal aminoacid sequence. Different aminoacid composition can also be invoked as one of the explanations, although different frequencies of *k*- mers with similar aminoacid composition prevent it from being considered the sole source of *k*-mer distribution. Some of the specificity can be the effect of translating DNA with a specific distribution of 3k-mers, which in turn is created by a range of DNA-specific factors such as GC-content, codon usage, presence of specific sites like splicing regulators and so on. If the analogy with codon usage bias is anything to go by, though, we should presume that there isn't a single source of selective pressure on *k*-mer composition. All the factors described above probably apply to some degree, as well as many others.

## Material and methods

CEGMA dataset of highly conserved genes from six model eukaryotic species (*A. thaliana, C. elegans, D. melanogaster, H. sapiens, S. cerevisiae, S. pombe*) was used. These are genes from 459 distinct orthogroups, each of which is represented by no more than one sequence from every species, for a total of 456-458 proteins per species [8].

50 randomly selected proteins from each species were used as a test set, and naïve Bayesian classifier (similar to multinomial classifier in [9]) was trained on the remaining ones. Test set sequences were assigned to the proteomes using this classifier using for values of *k* between 3 and 10. Similar procedure was performed on the complete proteomes of these species, using 10% of proteins randomly sampled as a testing set. All distances between *k*-mer distributions were calculated using FFP distance metric [10].

## Authors’ contributions

AM has conceived the analysis, performed it and written the paper.

## Acknowledgements

This work was supported by the Federal Agency of Scientific Organisations (Russian Federation) project #0345-2016-0031. Author is grateful to Dr. Y.P. Galachyants and A.N. Gurkov for the valuable discussion.

